# Loss of chromosome Y in primary tumors

**DOI:** 10.1101/2022.08.22.504831

**Authors:** Meifang Qi, Jiali Pang, Irene Mitsiades, Andrew A. Lane, Esther Rheinbay

## Abstract

It has long been recognized that certain cancer types afflict female and male patients disproportionately. The reasons for this disparity include differences in male/female physiology, effect of sex hormones, risk behavior and environmental exposures, and different copy number of the sex chromosomes X and Y. Loss of Y (LOY) is common in peripheral blood cells in aging men, and this phenomenon is associated with several diseases. However, the frequency and role of LOY in tumors is little understood. Here, we present a comprehensive catalog of LOY in >5,000 primary tumors from male patients in the TCGA. We show that LOY rates vary by tumor type, and provide evidence for LOY being either a passenger or driver event depending on context. LOY in uveal melanoma specifically is associated with age, survival and is an independent predictor of poor outcome. LOY creates common dependencies on *DDX3X* and *EIF1AX* in male cell lines, suggesting that LOY generates unique vulnerabilities that could be therapeutically exploited.

## Introduction

Gender and biological sex have been implicated in cancer incidence, mortality and response to therapy. These sex differences are caused by different physiology, the effect of sex hormones, environmental exposure and intrinsic genetic sex differences caused by the dichotomous sex chromosomes. Sex chromosome loss is the most frequent somatic change in peripheral blood lymphocytes of elderly individuals^1–4^ and sporadically occurs in normal tissues^5^. Specifically, loss of the Y chromosome (LOY) in aging men has been linked to increased mortality, incidence of hematologic and solid cancers, and Alzheimer’s disease^1–3^. Along with other copy number alterations, LOY has also been observed in some tumor types, including papillary renal cancer^6,7^, esophageal cancer^8^ and advanced prostate cancer^9^. The Y chromosome is comprised of a male-specific region (MSY), unique to this chromosome, and the pseudoautosomal regions (PARs), short genomic stretches near the ends of X and Y that undergo homologous recombination between the two sex chromosomes during meiosis. The MSY contains several ubiquitously expressed genes^10^, some of which are evolutionary conserved ancestral and potentially functional homologs of X inactivation escape genes^11^. Furthermore, somatic mutations in “escape from X-inactivation tumor suppressor genes (EXITS)” genes are enriched in tumors that also have loss of the second sex chromosome (X or Y)^12^. Little is known about the role of specific Y-linked genes in cancer, yet anecdotal evidence suggests relevance to disease biology. For example, aberrant expression of the Y gene *TSPY1* in females with dysgenetic gonads is associated with gonadoblastoma^13^. In prostate tumors, expression of the Y-linked histone demethylase *KDM5D* is associated with response to chemotherapy^14^; and *KDM5D* loss through LOY increases viability of renal cancer cell lines^15^.

Despite this important evidence for a role of Y in cancer, many whole-exome (WES) and whole-genome sequencing (WGS) cancer studies have nearly universally neglected analysis of the Y chromosome or deliberately excluded it, even in studies focused on cancer sex differences^16^. Reasons include the very small number of genes on Y that are relevant outside of male spermatogenesis and technical challenges caused by its haploid ground state, homology with X regions, and repetitive sequence from gene expansions. Prior studies on chromosome Y loss have focused on assessing the fraction of LOY cells in normal blood from SNP array^2,3^, evaluated loss of gene expression from Y^17^, or have been limited to fewer tumor types^12,18^. However, no comprehensive study of LOY in primary tumor tissue has been performed to date.

Our lack of knowledge around the Y chromosome in cancer creates missed therapeutic opportunities: LOY can expose specific cellular vulnerabilities caused by loss of genes without “backup copies”, those with homologs on X^12^,and loss of heterozygosity and potential haploinsufficiency of genes in the PARs.

We here present a comprehensive analysis of the Y chromosome across >5,000 male tumors from The Cancer Genome Atlas (TCGA), suggesting both driver and passenger roles for LOY in different cancer types.

## Results

### LOY is a frequent somatic event in primary cancers

Many commonly used somatic copy number calling methods do not provide faithful copy number estimates for the Y chromosome. We therefore adapted an established copy number analysis method^19^ to analyze X, Y and the two pseudoautosomal regions (PAR1 and PAR2) separately (**Figure 1A;** see Methods for details) in 5014 male and 5394 female primary tumors using whole exome sequencing (WES) from The Cancer Genome Atlas (TCGA; **Supplementary Table 1,2**). Additionally, we derived a method based on Y gene expression to measure “functional” LOY (fLOY) from RNA-Seq as an alternative method for samples without WES or normal control data, confirming that gene expression can reliably detect LOY^17^ (**Figure 1A**; see Methods for details). We observed a range of Y copy events, most frequently complete LOY (28%). Relative LOY in the presence of additional copies (rLOY) also occurred at 2% frequency (Methods) (**Figure 1B, C**). Overall, 1504 of 5014 male tumors harbored either complete or relative LOY. As expected, LOY calls from exome were accompanied by loss of expression of Y-linked genes^10^ (**Figure 1B, D**) and a median reduction of PAR gene expression by 32% compared to wild-type Y samples. Indeed, fLOY from TCGA tumor RNA-seq alone gave nearly identical results with most differences for lower-purity tumors with considerable normal cell content **(Figure 1B, Supplementary Figure 1, 2; Supplementary Table 1**). For comparison, loss of the entire X chromosome (LOX) was always relative, consistent with essentiality of X for cellular survival (**Supplementary Table 2**). LOX in females was accompanied by loss of *XIST* expression, which induces X inactivation when more than one X chromosome is present and a 26% decrease of PAR gene expression (**Supplementary Figure 3A**). In total, 757 of 5394 (14%) female tumors experienced LOX. Interestingly, LOY rate was significantly higher in Black (39%) compared to Asian (29%) and white (28%) participants (Proportion test *P*=8.3 x 10-5 and *P*=0.006; **Supplementary Figure 4A**) but no difference was observed for LOX (*P*=0.21 and *P*=1; **Supplementary Figure 4B**). Because of its frequency and the potential impact of losing an entire chromosome without a backup copy, we focused on complete LOY for further analyses.

**Figure 1:**
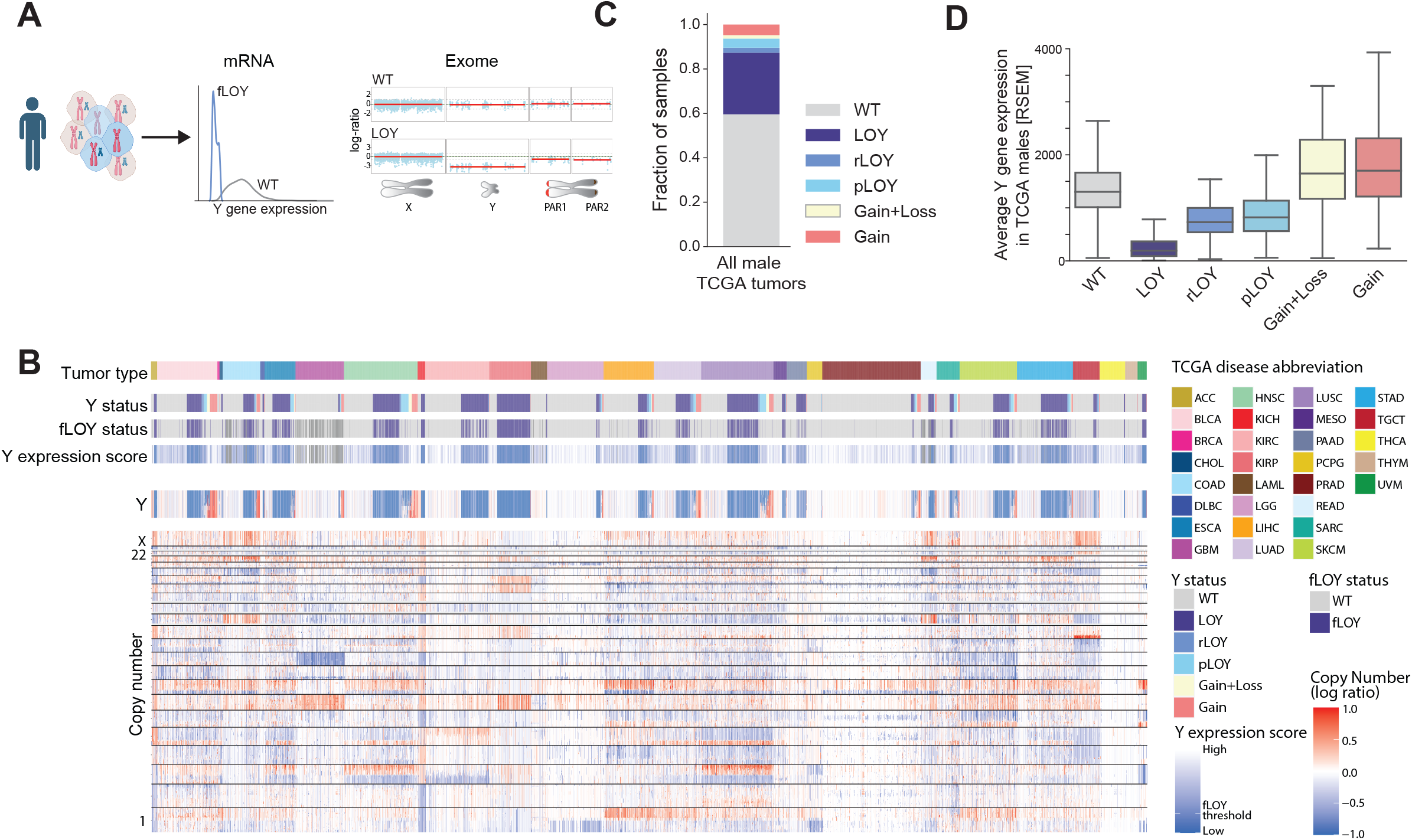
Schematic of methods to detect LOY in tumors. LOY can be detected from tumor gene expression profiles (mRNA-Seq or microarray) or exome (WES). fLOY: “functional LOY (expression-based). **B**, Copy number profiles of 5014 male tumors from TCGA with the Y chromosome copy number shown disproportionally large. Y gene expression score, expression-based loss calls (fLOY) and Y status inferred from exome copy number are shown. Tumor types are labeled with TCGA tumor codes. **C,** Fraction of tumors with indicated Y chromosome alterations among male TCGA tumors. pLOY: partial LOY; rLOY: relative LOY (total Y copy number ≥1); LOY: total Y copy number equals zero. Details are described in the Methods. **D**, Average Y gene expression for each type of Y alteration corresponds to inferred copy state.

### LOY rates across tumor types

Like other patterns of chromosomal gains and losses, rates of different Y chromosome events varied by cancer type (**Figure 2A; Supplementary Table 3)**. Complete LOY was most frequent in renal papillary cancer (KIRP; 80%) and esophageal cancer (ESCA; 57%), and as low as <2% in pheochromocytoma and paraganglioma (PCPG) and thymoma (THYM), and these fractions were consistent with previously published results based on cytogenetics or WGS^6–8,18,20^. Statistical evaluation of arm-level losses with GISTIC (Methods) confirmed significant loss of Yq and Yp in many tumor types (**Figure 2B, Supplementary Table 4**). In contrast, LOX was most frequent in KICH (56%) and uveal melanoma (UVM; 43%) and but near absent in thyroid cancer (THCA; 1%) and THYM (0%) (**Supplementary Figure 3B; Supplementary Table 3**). In general, LOY occurred much more frequently than LOX, with major differences in KIRP and ESCA, suggesting a specific selective advantage of LOY in these tumor types (**Supplementary Figure 3C**). LOY and LOX rates were nearly equally frequent (>20%) in KICH, UVM, MESO and SKCM (**Supplementary Figure 3C**), possibly because loss of a common or homologous gene on X and Y might drive selection of tumors with sex chromosome loss.

**Figure 2:**
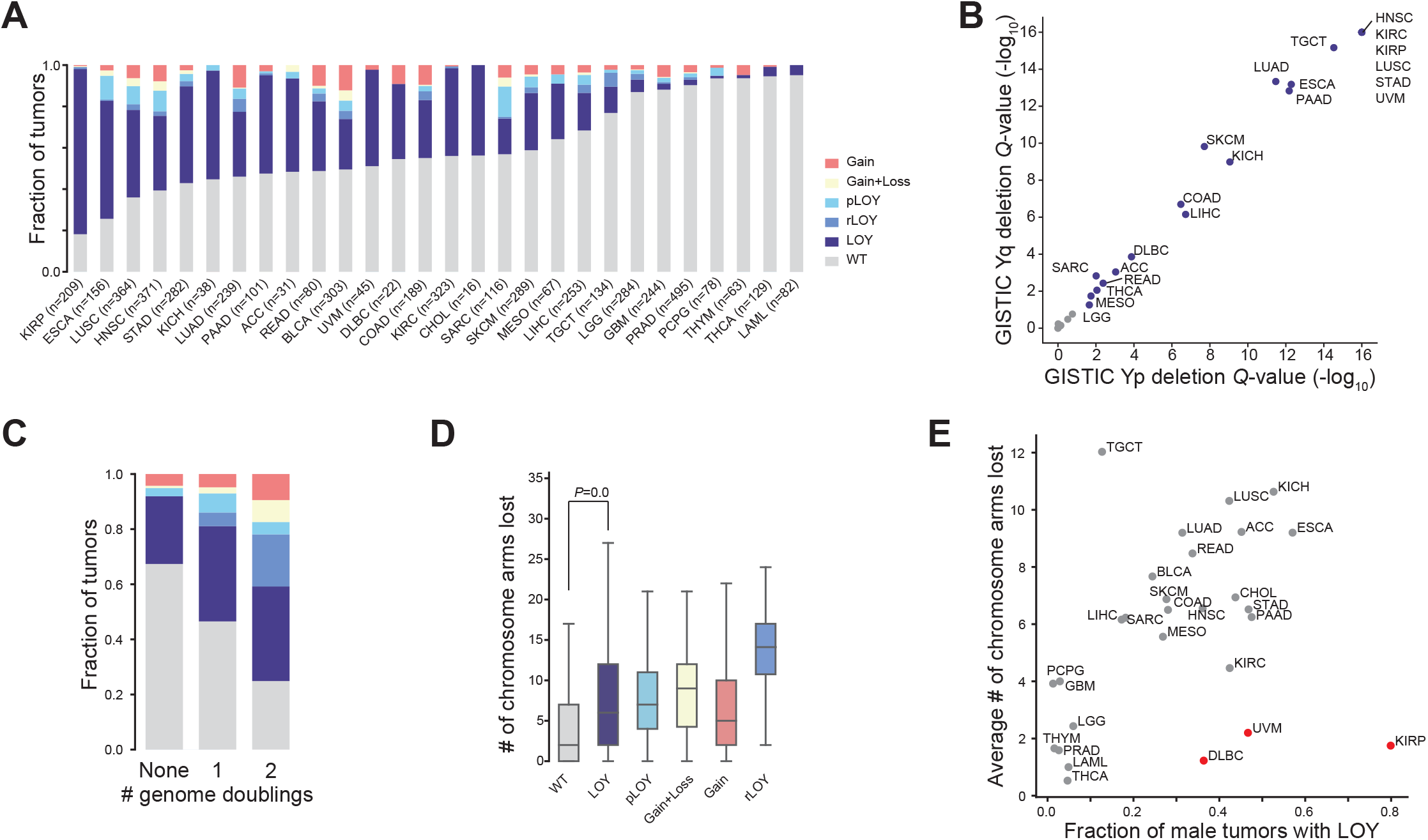
Fraction of different Y chromosome alteration for each tumor type. **B,** GISTIC *Q*-values for Yp and Yq chromosome loss confirm recurrent losses in many tumor types. Strong correlation between Yp and Yq further suggests that Y is typically lost as whole chromosome. **C**, Fraction of Y alterations in tumors with no, one or two genome doublings. **D**, Number of autosomal chromosome arms lost in tumors with WT, Y loss or other Y alterations. **E**, Correlation between total arm losses (autosomes only) and LOY. Red dots indicate tumor types with high LOY rate and low overall arm-loss rate.

### LOY is common in aneuploid tumors

As a small, gene-poor chromosome, Y has a relatively high chance of being lost from cells “by chance”^21^, and the paucity of genes on this chromosome that are expressed outside of the male reproductive system suggests that there could be little selective pressure for its retention. In peripheral blood, somatic LOY has been associated with generally increased genomic instability^3^. To answer whether somatic LOY is correlated with genomic instability in primary tumors, we compared LOY rates with genome doubling events and aneuploidy, measured as arm-level losses of only the autosomes^22^. Concordant with unstable genomes, tumors that had undergone genome doublings were more likely to harbor various Y copy changes (**Figure 2C**). LOY tumors also had significantly more arm-level losses (median 6, IQR 2-12) compared to WT tumors (median 2, IQR 0-7; MWU *P*=0.0, **Figure 2D**). As expected, loss rates were highest in tumors with rLOY as a product of genome doubling (**Figure 2C, D**). On a tumor-type level, LOY rates were weakly correlated with mean arm loss scores (Pearson’s *r*=0.33; *P*=0.08) (**Figure 2E**), suggesting that LOY follows general aneuploidy rates. UVM, KIRP and diffuse large B-cell lymphoma (DLBC) had a comparatively high fraction of LOY tumors (**Figure 2E**), suggesting selection for LOY in specific diseases.

### Association of LOY with point mutation drivers

We next assessed whether LOY was associated with mutations in cancer driver genes^23^. The most significantly associated driver across the pan-cancer cohort was *TP53* (Fisher’s Exact *P*= 1.23E-46), the most frequently altered tumor suppressor in cancer whose loss causes genomic instability (**Figure 3A**; **Supplementary Table 5**) and confirming the link to aneuploidy. In total, 47.1% of LOY tumors had a damaging somatic mutation in *TP53*, compared to only 25.7% of tumors with intact Y (**Figure 3B**). In addition, we found *KDM5C* and *KDM6A* mutations enriched in the LOY pan-cancer cohort, and in individual tumor types (*KDM5C* in KIRC with *P*=1.09E-4 and *KDM6A* in BLCA with *P*=6.09E-3 and LUSC with *P*=0.034; **Figure 3A, Supplementary Table 5**)^12,15^. Both genes have homologs on Y, indicative of a two-hit event^12^.

**Figure 3:**
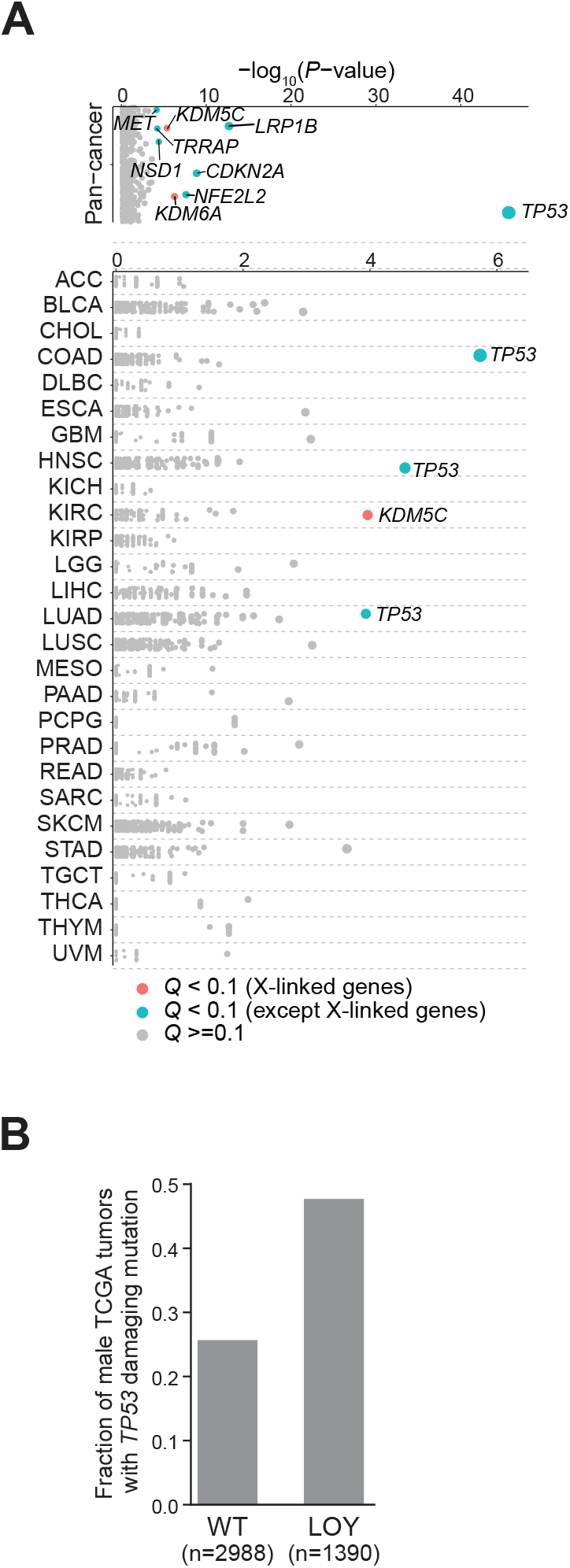
Cancer genes with somatic mutations enriched in LOY compared to WT male tumors. Top panel: Pan-cancer; bottom panel: individual tumor types. X-axis indicates Fisher’s test *P-* value. Genes with *Q*-value <0.1 are highlighted. **B,** Fraction of male LOY and WT tumors with *TP53* damaging mutation.

### Driver candidates on the Y chromosome

To identify individual candidate tumor suppressors on Y, we queried our catalog of Y copy number calls for recurrent focal deletions in tumors without LOY (Methods). Only two tumor types, head and neck and lung squamous carcinoma (HNSC and LUSC), had a focal deletion on Yq potentially extending into the heterochromatin region (**Supplementary Table 6).** In LUSC, this region includes the ubiquitously expressed^10^ tumor suppressor *KDM5D*^9,15^ and *EIF1AY*, homolog of the X-linked cancer gene *EIF1AX.* All focal deletions occurred in tumors with many copy number changes, thus further functional study will be required to determine whether *KDM5D* and/or *EIF1AY* are tumor suppressors in LUSC.

We also searched for MSY and PAR genes with recurrent point mutations (SNVs, short indels) in tumors without LOY as potential tumor suppressors (Methods), reasoning that they might be inactivated by point mutations in the absence of LOY. However, no Y-linked genes were significantly recurrently altered (**Supplementary Table 7**). A caveat to this analysis is the careful filtering of the TCGA pan-cancer mutation calls, including panels-of-normals and read depth requirements^24^, could lead to overconservative removal of mutations on the Y chromosome.

### LOY predicts patient outcome

To test whether somatic LOY is associated with patient outcome, we performed survival analysis for all male TCGA cases with progression-free survival (PFS) and chromosome Y status information. Although a pan-cancer analysis is necessarily influenced by tumor type, we observed a poorer outcome in tumors with LOY (**Figure 4A;** hazard ratio 1.17, log-rank *P*-value <0.00117). At the individual tumor type level, we found that LOY was significantly (Q<0.05) associated with poor PFS in three tumor types: uveal melanoma (UVM; HR=6.4, log-likelihood ratio test *P*=0.0016, *Q*=0.0165), prostate adenocarcinoma (PRAD; HR=3.7, *P*=0.001, *Q*=0.0165) and mesothelioma (MESO; HR=2.7, *P*=0.002, *Q*=0.0165) (**Figure 4B, C; Supplementary Table 8).**

**Figure 4:**
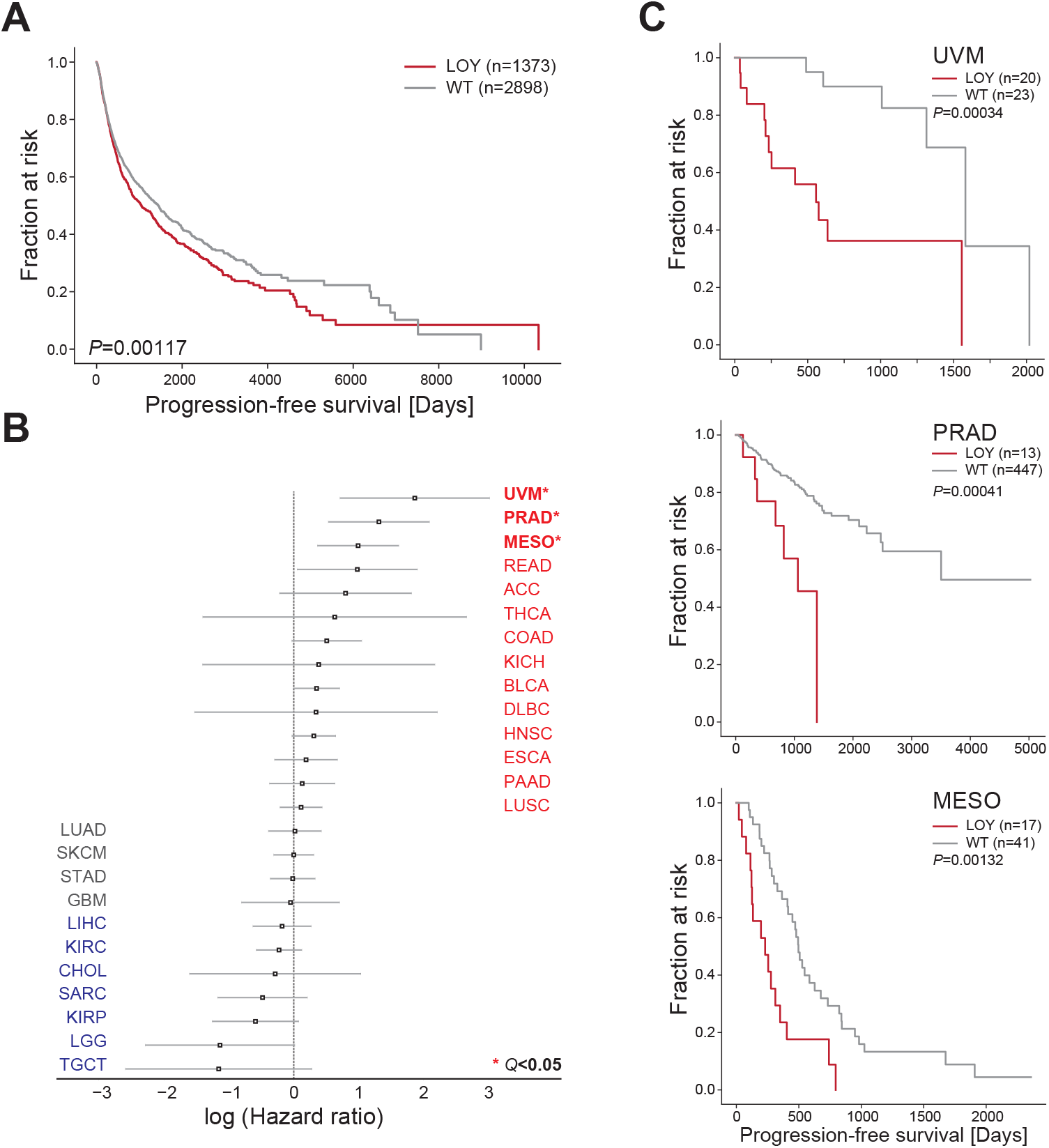
Kaplan-Meier survival statistic depicts progression-free survival for male tumors with complete LOY compared to WT tumors. **B**, Hazard ratios (log) and 95% confidence intervals for LOY for each tumor type, sorted by hazard ratio. Bold and * indicate significant tumor types (*Q*<0.1). **C,** Kaplan-Meier survival curves for tumor types with significant contribution of LOY to survival. UVM: uveal melanoma; PRAD: prostate adenocarcinoma; MESO: mesothelioma.

### LOY is a candidate driver event in uveal melanoma

The strong difference in PFS and clonality of LOY in uveal melanoma (**Figure 4B, C**) prompted us to further study the role of LOY in this tumor type. Uveal melanoma is a rare malignancy of melanocytes in the eye. In contrast to melanoma of the skin, this disease is not associated with ultraviolet light exposure and its defining genomic aberrations differ from the classic cutaneous melanoma events^25,26^. This includes an overall stable genome with well-characterized broad copy number changes and no *TP53* mutations^26,27^. LOY was observed in uveal melanoma decades ago, but not investigated much further^28^. For unknown reasons, uveal melanoma incidence is biased towards males, and this bias increases with patient age^25^ (**Figure 5A**). Interestingly, we found a 10-year difference in median age at diagnosis between WT and LOY for uveal melanoma in the TCGA cohort (median 58 vs 68, MWU *P*=0.006; **Figure 5B**), prompting the question whether age-related or environmentally induced LOY occurs in uveal melanocytes and whether LOY is a driver event in this disease. Notably, the difference in outcome was not explained by age (*P*_LOY_=0.01; *P*_age_=0.32; Methods).

**Figure 5:**
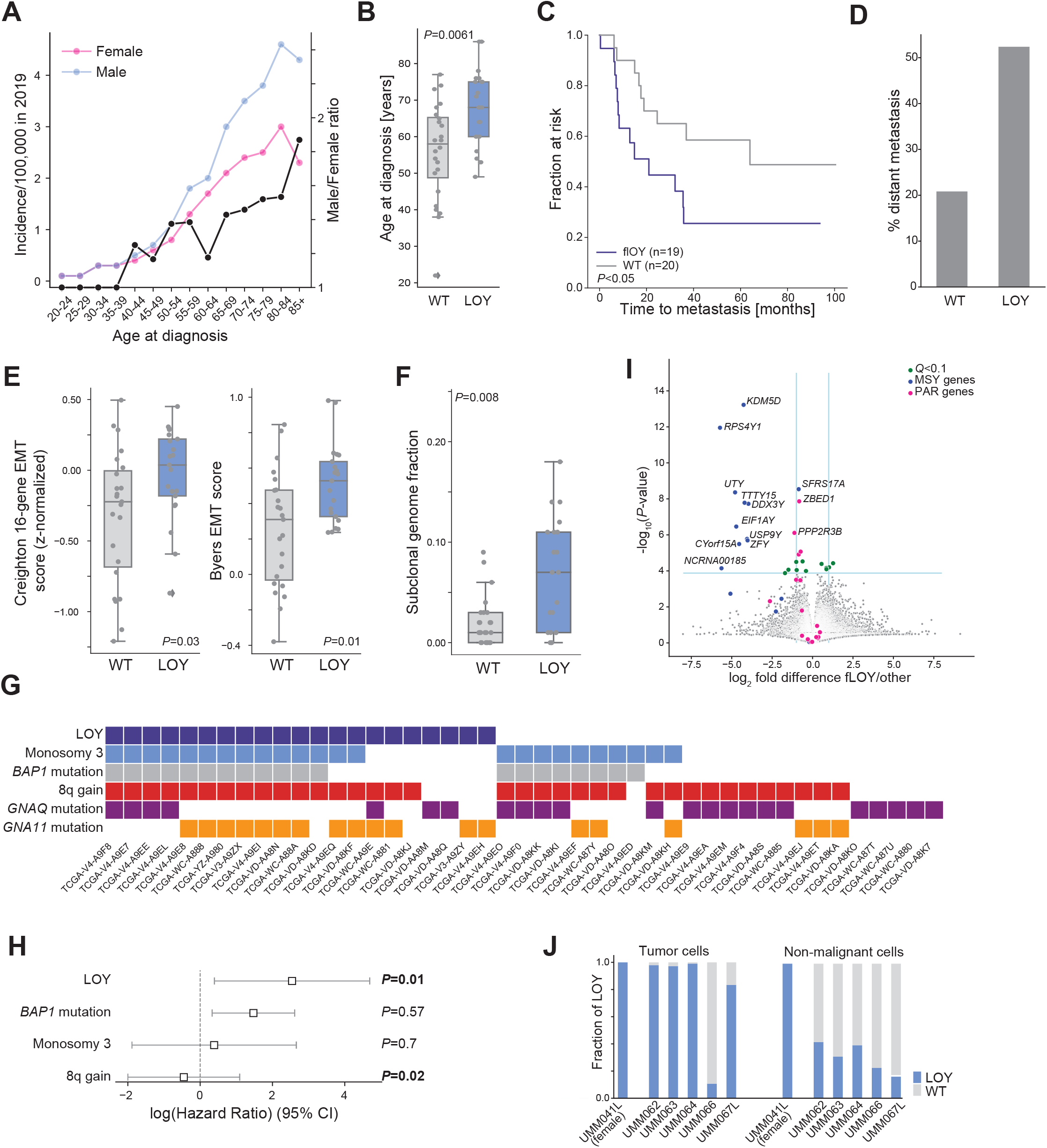
SEER incidence for uveal melanoma for male (blue) and female (pink) patients. The male/female incidence ratio is shown in black (secondary Y-axis). **B,** Distributions of age at diagnosis for LOY and WT male uveal melanoma patients from TCGA. **C**, Kaplan-Meier survival curve for male patients from an independent cohort^45^. **D,** Percentage of TCGA patients with distant metastasis with LOY or WT. **E**, Distribution of epithelial-to-mesenchymal (EMT) scores^30,31^ for LOY and WT TCGA tumors. **F**, Distribution of subclonal genome fraction^26^ for TCGA tumors. **G**, Somatic mutations and LOY for each TCGA uveal melanoma tumor. **H,** Cox proportional hazards for known predictors of poor survival in uveal melanoma and LOY. **I,** Volcano plot showing differentially expressed genes between LOY and WT uveal melanoma in TCGA. MSY: male-specific region on Y; PAR: pseudoautosomal region. **J,** Fraction of tumor cells (left) and non-malignant single cells^33^ (right) with LOY (blue) or intact Y (WT, grey). One female patient (UMM041L) is shown as control.

We validated the remarkable difference in outcome between LOY and WT tumors in an independent cohort of 39 male uveal melanoma patients with gene expression profiles^29^ (Methods; **Supplementary Figure 5A**). Confirming our prior results, 49% (19/39) of these patients had Y expression based, “functional” LOY (fLOY) (compared to 47% in the TCGA WES cohort) and outcome for these patients was significantly worse (**Figure 5C;** *P*=0.045). Recapitulating the age difference in the TCGA cohort, age at diagnosis was significantly higher for patients with fLOY tumors (68 vs. 60 years; *P*=0.01).

Metastatic spread is common in patients with uveal melanoma, and once a tumor has metastasized, prognosis is very poor^25^. We found that LOY tumors were significantly more likely to metastasize than wild-type tumors (52% vs 21%, *P*=0.03, Fisher’s Exact test), and this trend was confirmed in the validation data set (68% vs 45%, *P*=0.2) (**Figure 5D; Supplementary Figure 5B**). In addition, epithelial-to-mesenchymal (EMT) gene expression scores^30,31^ (**Figure 5E**) and subclonal fraction^26^ (**Figure 5F**), associated with metastatic potential and aggressiveness of cancer cells, were significantly higher in LOY compared to WT cases (Mann-Whitney U *P*<0.03 for EMT; subclonal fraction Mann-Whitney U *P*=0.008).

Uveal melanomas are characterized by ubiquitous early mutations in *GNAQ/GNA11*, monosomy of chromosome 3, 8q gain, mutations in and loss of the *BAP1* tumor suppressor and mutations in the *EIF1AX* elongation and the *SF3B1* splicing factors^26^. Monosomy 3, *BAP1* (located on chromosome 3) and 8q have been linked to prognosis, while *EIF1AX* predicts better outcome. To test whether the LOY outcome difference seen in the two cohorts could be explained by these underlying, well-described alterations, we calculated the overlap of events across tumors (**Figure 5G**). There was no association between *BAP1* alterations and LOY (Fisher’s Exact *P*=0.14), chr3 monosomy (*P*=0.14) or 8q gain (*P*=0.73). A multivariate Cox proportional hazard model that accounted for these bad outcome predictors maintained LOY as the most significant predictor (**Figure 5H;** *P*=0.01). Importantly, we observed no significant overlap of *EIF1AX* mutations and loss of its Y-linked homolog *EIF1AY* through LOY (*P*=0.61), consistent with their opposite predictive effects on patient outcome. Interestingly, *GNA11* mutations were enriched in LOY tumors (*P*=0.017), suggesting a potential selective advantage of LOY in the presence of the *GNA11* but not *GNAQ* genotype. Aside from a single *DDX3Y* missense mutation, no mutations in Y-linked genes were detected in the UVM TCGA cohort^26,32^.

Lack of somatic point mutations on Y and the fact that Y is lost in its entirety in uveal melanoma complicate the identification of a single tumor suppressor. We therefore compared gene expression profiles between male LOY and WT tumors to identify differentially regulated candidate Y-linked drivers. As expected, nearly all significantly differentially expressed (DE) genes are located on Y (either in the MSY or PAR; **Figure 5I; Supplementary Table 9).** Among DE genes, chromatin regulators *KDM5D* and *UTY (KDM6C)* are among the most significantly downregulated in LOY tumors (**Supplementary Figure 5C).** *KDM5D* demethylates the active histone mark H3K4me3. Loss of *KDM5D* interferes with regulation of normal gene expression and has been shown to accelerate cell cycling^9^. This is particularly intriguing in the context of frequent mutations and loss of the histone ubiquitinase *BAP1*, suggesting multifaceted alterations of chromatin regulation in uveal melanoma. One potential driver function of LOY might thus be the loss of the *KDM5D* tumor suppressor.

Next, we sought to understand the clonality of LOY in uveal melanoma by directly assessing the fraction of functional LOY in single cell gene expression profiles^33^ (Methods). Four out of five male tumors had substantial to complete (83 to >99%) fLOY, with much smaller percentages of fLOY in non-malignant cells (17-42%; **Figure 5J**). Single-cell gene expression experiments can suffer from gene detection limits (“drop out”), which likely contributes to the high fLOY rates in non-malignant cells. Yet, fLOY cells were not of lower quality overall than WT cells (**Supplementary Figure 5D**). The large number of tumor cells with fLOY among single cells suggests that loss of the Y chromosome is an early, clonal event in this disease.

Our sex chromosome calls suggest that in addition to LOY in males, LOX is also common in female uveal melanoma with nearly equal frequency (43%; **Supplementary Figure 3B**). Similar to LOY tumors, LOX cases had increased rates of metastasis (47% vs 15%, Fisher’s Exact *P*=0.06; **Supplementary Figure 5E**). In contrast to LOY tumors, there was no significant age difference (MWU *P*=0.8). These results suggest that sex chromosome loss is not simply a consequence of age, and implies genetic driver events among shared homologous genes on X and Y.

### Differential dependencies in LOY cell lines

Finally, we searched for gene dependencies that are found in male cell lines with LOY as specific, potentially targetable vulnerabilities^34^. Top dependencies among fLOY lines in aggregate (**Figure 6A; Supplementary Table 10;** Methods) were *DDX3X* and *EIF1AX*, X-linked homologs of *DDX3Y* and *EIF1AY. DDX3X/DDX3Y* are RNA helicases implicated in transcription. *EIF1AX/EIF1AY* are essential and ubiquitously expressed translation initiation factors, and *EIF1AX* is somatically mutated in several cancer types^32^. Concordantly, Y-linked *DDX3Y* and *EIF1AY* were among the top four differentially expressed genes between LOY and WT cell lines (**Supplementary figure 6).** It is important to note that CRISPR gene effect scores suggest the dependency on these two genes is exclusive to LOY cell lines without dependency in lines with intact Y (crossing the dependency threshold of −1; **Figure 6B).** This includes a uveal melanoma cell line, Mel270, that is strongly dependent on these two genes (gene effect scores −1.195755 and −1.326149, respectively). *DDX3X* and *EIF1AX* thus represent two critical vulnerabilities specific to LOY cells that could be therapeutically exploited.

**Figure 6:**
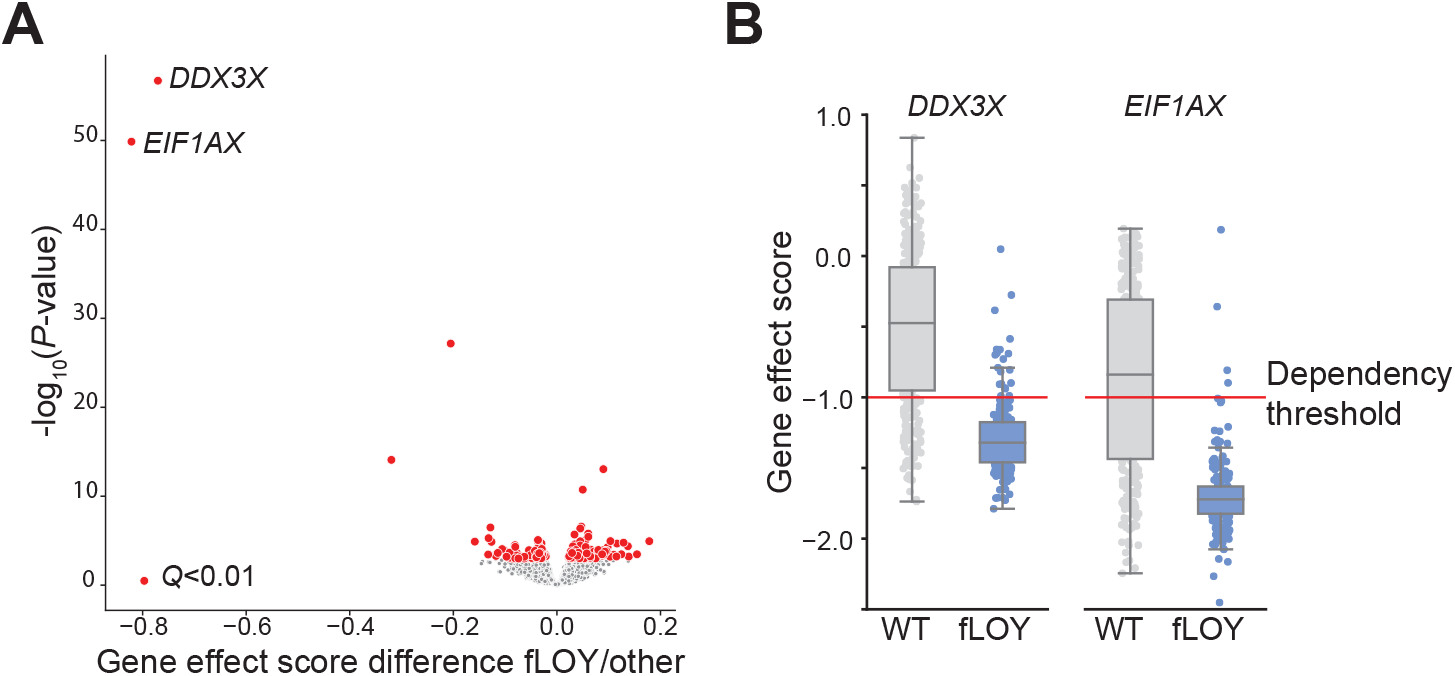
Volcano plot depicting differential dependency for each gene in LOY vs. WT cell lines from CCLE^34^. Red dots indicate significantly different dependencies. **B**, Gene effect scores for *DDX3X* and *EIF1AX* show no dependency on these genes in cell lines with Y, and strong dependency in LOY lines. Dependency map-defined threshold for dependency (−1) is shown as red line.

## Discussion

Anecdotal evidence has suggested that chromosome Y might play a driver role in some cancer types. Through development of specific methods to analyze copy number of chromosome Y from genomic and transcriptomic profiles, our study presents the first comprehensive analysis of Y copy number variation across 29 tumor types of different lineages. We show that LOY is extremely common in many tumor types, with frequencies higher than some classic driver genes. LOY can occur in the context of overall genomic instability, where LOY is likely a passenger. However, an additional contribution to tumor biology even in the case of a passenger event remains possible, such as if LOY creates a paralog dependency^35^, and warrants further study. We also present evidence for a driver role of LOY in uveal melanoma, where LOY is a clonal event that is strongly associated with predictors of poor survival, including older patient age, tumor heterogeneity, proliferation, and time to progression. Yet LOY is independent and potentially more predictive than other well-known somatic alterations in uveal melanoma, suggesting it as a prognostic biomarker that is relatively easy to detect through standard clinical assays.

Because chromosome Y usually does not have a “backup” copy, LOY leads to loss of several unique and ubiquitously expressed genes, with potentially important implications for cell fitness (e.g. *KDM5D, KDM6A, RPS4Y1).* In addition, loss of a copy of the PARs introduces LOH and could cause decreased expression of several immune genes, including the macrophage receptors *CSFRA* and *IL3RA* and the immune surface gene *CD99. CD99* is highly expressed in glioblastoma cells, where it correlates with migration and invasiveness^36,37^. The dependence of GBM tumors on *CD99* could explain the relatively low LOY rates observed in this disease. LOY creates unique and targetable vulnerabilities through loss of these genes and exposing of potentially altered X-linked genes.

A limitation of our study is the lack of evidence for recurrent alteration of a specific tumor suppressor on Y, as could be assessed by focal deletion or recurrent somatic mutation. This might be caused by lack of sensitivity of our methods (especially point mutation calling). It is also possible that LOY creates a unique fitness advantage through loss of multiple tumor suppressors simultaneously. Further functional studies will be required to complement the existing literature on Y-linked tumor suppressors across all tumor types with frequent LOY.

## Methods

### TCGA data

This study used TCGA Pancancer Atlas obtained from https://gdc.cancer.gov/about-data/publications/pancanatlas. Participant information was retrieved from file TCGA-CDR-SupplementalTableS1.xlsx and purity/ploidy were from TCGA_mastercalls.abs_tables_JSedit.fixed.txt.

### Somatic copy number

We used the FACETs copy number caller^19^ framework as a base for calling Y copy number changes, with several modifications: (i) In the human hg19 genome assembly, the two PAR regions at the ends of the X and Y chromosomes are exclusively assigned to the X chromosome, leading to discordant X copy number and heterozygous SNP calls in these regions in tumors with XY genotype. We therefore separated the PARs from X and treated them as additional autosomes. For simplicity, we assume that in normal control tissue of most males, there will be exactly one copy of each X and Y, and two X and zero Y copies in the majority of female tissues (see below for limitations). All autosomes will typically have two copies regardless of biological sex. (ii) The Y chromosome contains a large heterochromatin region as well as repetitive stretches of sequence that can lead to incorrect copy calls. We therefore identified ambiguously mappable regions on Y by counting read depth in contiguous 500 bp bins in female TCGA normal samples and two female WGS samples. Bins in which at least one position had a read depth ≥ 10 in WGS samples were excluded from analysis. We also excluded positions that had at least read depth ≥ 10 in ≥ 20% female samples from TCGA. Additionally, we excluded regions on the X and Y chromosomes with low mappability^38^ (score <0.5); and ubiquitously high coverage^38^ and the Y centromere^39^ (from GATK). (iii) We used the dbSNP151 set of common germline variants to gather allelic counts on all chromosomes. To increase the number of coverage data points on gene-poor Y, we added additional pseudo SNPs (“pseudo_snps=100”) and coverage count at the middle position for each exon of chromosomes X and Y to the coverage pileup file. (iv) Because the segmentation parameter cval is sensitive to the number of data points (SNPs), we adjusted cval=50 to increase the copy segment resolution on Y while leaving the default cval=150 for all other chromosomes.

With these modifications, we used the FACETs framework to estimate total copy number (tcn) for Y. In cases where purity and ploidy estimates diverged substantially (ploidy difference ≥ 1 or purity difference ≥ 0.2) from previously published TCGA values (TCGA pan-cancer file TCGA_mastercalls.abs_tables_Jsedit.fixed.txt), we corrected tcn with published TCGA purity and ploidy values. Official TCGA calls are based on the ABSOLUTE method^40^, which relies on copy number as well as somatic mutations as data source; this is advantageous especially for tumors with few copy alterations where purity and ploidy estimates from copy number alone can be less reliable. The correction was calculated as

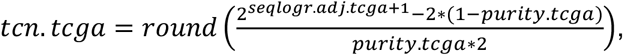

Where *cnlr.median* is the median log ratio, *seqlogr. adj. tcga = cnlr. median – digLogR.tcga* and 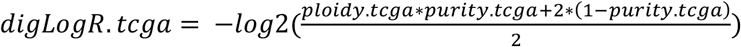.

The expected maximum Y WT copy number (max.Y.WT) is defined as round(ploidy/2) if round(ploidy) is even or round(ploidy/2+1) if round(ploidy) is odd. The minimum Y WT copy number (min.Y.WT) is defined as round(ploidy/2) if round(ploidy) is even or round(ploidy/2-1) if round(ploidy) is odd. We call a segment is “gained” if tcn>max.Y.WT and a segment “lost” if tcn<min.Y.WT. A segment on Y is considered “WT” if min.Y.WT ≤ tcn ≤ max.Y.WT. Then we classified CNVs of Y chromosome into several groups (**Supplementary Figure 7**): (1) WT: All segments on Y are “WT”; (2) Gain: At least one segment on Y is gained and potential other segments are WT; (3) Gain+Loss: at least one segment is gained and at least one segment is lost; (4) pLOY (partial LOY): at least one segment on Y is lost, and all others are WT; (5) rLOY: All segments of the Y (≥99% of the Y chromosome) are lost, and tcn ≥ 1 (e.g. after genome doubling) (6) LOY: All segments on Y (≥99% of the Y chromosome) are lost, and tcn=0.

X chromosome status classification is calculated similarly, except that LOX is compared with round(ploidy), and LOX is defined as ≥99% of the X chromosome is lost (**Supplementary Figure 7**). We identified 39 tumors with discordant reported gender and genomic sex or low quality by manually review, and these were not included in further analysis (**Supplementary Table 1, 2**). Recurrently mutated broad and focal copy number changes on X and Y were identified with GISTIC2^41^ with parameter -rx 0 to include the sex chromosomes. BLCA was excluded from Figure 2B due to lack of markers to evaluate Yq. Yq peak regions were manually trimmed back to the boundary with the heterochromatin region after automatic extension with markers by GISTIC to limit the focal peak to regions with measured coverage data points. Significant Y regions were annotated with the Gencode V40 gene list^42^.

A caveat of our analysis is that LOY rates are likely conservative and underestimated for two reasons: (i) most controls in TCGA are peripheral blood samples that may be subject to age-related somatic LOY, and thus tumor Y copies can be “gained” relative to control; (ii) if the fraction of somatic LOY in the control and tumor are approximately equal, no difference will be detected and LOY is not called.

### Classification of LOY based on gene expression (fLOY)

Gene expression RSEM values were downloaded from the GDC Pancancer Atlas website (File EBPlusPlusAdjustPANCAN_IlluminaHiSeq_RNASeqV2.geneExp.tsv). A list of genes in the male specific region of Y was used to identify a seven-gene signature of genes consistently expressed in normal tissues to score expression from this chromosome *(RPS4Y1, DDX3Y, KDM5D, USP9Y, EIF1AY, UTY, ZFY;* **Supplementary Figure 1A, B**). A common set of housekeeping genes^43^ was used to control for overall expression activity. The ratio of the mean of expression of the seven Y genes to the expression across housekeeping genes was calculated as “Y expression score” for each individual sample. A ratio threshold of 0.035, corresponding to Y gene expression of 3.5% of housekeeping gene expression, was manually identified to classify tumors into functional” LOY (flOY) (ratio < 0.035) and “WT” (ratio ≥ 0.035) (**Supplementary Figure 1C, D**). 478 male patients did not have RNA-seq expression calls (mostly GBM) and were not included in the classification or downstream analysis based on Y status. Differential expression p-values were calculated with a *t*-test followed by the Benjamini-Hochberg FDR correction.

### Survival analysis

Survival analyses were conducted using the lifelines Python package (https://github.com/CamDavidsonPilon/lifelines/). Progression-free survival from the Pancancer Atlas (TCGA-CDR-SupplementalTableS1.xlsx) was used as endpoint. Significance for Kaplan-Meier statistics were calculated with the log-rank test. Hazard ratios were calculated with the Cox proportional hazards model and significance was assessed with the log-likelihood ratio test. Uveal melanoma analysis accounting for age included TCGA metric “Age at Diagnosis”. Only tumor types with at least five samples in either LOY or WT group were included.

### Genomic instability

Aneuploidy scores for TCGA cases were obtained from Taylor et al.^22^.

### Point mutations

TCGA Pancancer multi-center somatic mutation calls (mc3.v0.2.8.PUBLIC.maf.gz) were used for mutation analyses. Only variants with the following damaging classifications were included: ‘Frame_Shift_Del’, ‘Frame_Shift_Ins’, ‘In_Frame_Del’, ‘Missense_Mutation’, ‘Nonsense_Mutation’, ‘Splice_Site’, ‘Translation_Start_Site’. For assessing whether LOY was associated with mutations in known driver genes^23^, we used Fisher’s Exact test to compare mutations between the LOY and WT groups. Recurrence analysis for mutated Y chromosome genes was performed with MutSig2CV with default settings on only male cases without LOY and rLOY chromosome status.

### Cell line dependencies

Gene expression and CRISPR gene effect scores were obtained from the DepMap portal (https://depmap.org/portal/; version 22Q2). Cell lines were classified with respect to sex and fLOY status by using a combination of Y gene expression of the 7-gene signature described above and *XIST* expression levels. We excluded 12 annotated male cell lines with absent Y expression and high *XIST* expression as potentially female lines (although it is possible that some of these lines are male with LOY and X multisomy, where additional X chromosomes will undergo silencing. Differential gene expression and dependency for all cell lines in aggregate and each disease type were evaluated with a two-sided *t*-test followed by FDR multiple hypothesis correction.

### Incidence data for uveal melanoma

Incidence data for cases in the United States in 2019 for cancer site “eye and orbit” were downloaded from the NCI’s Surveillance, Epidemiology and End Results website (https://seer.cancer.gov/statistics-network/explorer/application.html) on May 11, 2022.

### Uveal melanoma validation cohort

Functional LOY scores were calculated with Affymetrix array gene expression data using the same seven Y-linked and housekeeping gene sets for 39 uveal melanoma samples from Laurent et al^29^. Classification into fLOY and WT tumors was obtained from the bimodal distributions of Y gene expression (**Supplementary figure 5A)** and survival analysis was performed as described above.

### Classification of single cells

Single-cell expression data from uveal melanoma tumors was obtained from Durante et al^33^. scRNA-seq data were processed and analyzed using R (4.0.5) Seurat package (4.0.1)^44^. Raw counts of 11 samples were read into R using the Read10X function and aggregated into one Seurat object. Several metrics were used to account for dead cells and droplets: 1) The number of unique molecular identifiers (UMIs) per cell; 2) The number of detected genes per cell; 3) The proportion of mitochondrial genes; 4) Number of genes detected per UMI (log_10_(number of detected genes) / log_10_(number of UMIs)). Only cells with UMIs greater than 500, expressed 250 and 8000 genes inclusive, had mitochondrial content less than 10% and >0.8 detected genes/UMI were retained for future analysis. After filtering, 52,294 cells were left. Data were normalized using the NormalizeData function in Seurat with LogNormalize setting and a scaling factor of 10,000. Principal component analysis (PCA) was used to reduce dimensionality with number of variable features set to 2000. Clustering was conducted with FindClusters using the first 20 principal components and 1.5 as resolution parameter. The original Louvain algorithm was utilized for modularity optimization, which resulted in 46 clusters. Identified clusters were visualized using t-distributed stochastic neighbor embedding (t-SNE) and they were annotated as described in the source using the following markers^33^: Tumor cells *(MLANA, MITF, DCT)*, T Cells (*CD3D, CD3E, CD8A*), B cells (*CD19, CD79A, MS4A1*), plasma cells (*IGHG1, MZB1, SDC1, CD79A*), monocytes and macrophages (*CD68, CD163, CD14*), NK Cells (*FGFBP2, FCG3RA, CX3CR1*), retinal pigment epithelium (*RPE65*), photoreceptor cells (*RCVRN*), and endothelial cells (*PECAM1, VWF*). To measure functional LOY, we calculated average Y expression for the seven most expressed genes (*DDX3Y, EIF1AY, KDM5D, RPS4Y1, USP9Y, UTY, ZFY)* and determined a threshold from female cell values as

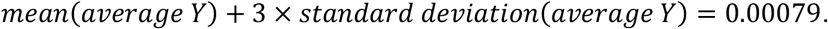

Cells with average Y expression smaller than the threshold were classified as LOY, “WT” otherwise.

### Statistics

Non-parametric comparisons were performed using the Mann-Whitney-U test implemented in the Python scipy.stats package. Plots were generated in Python using the Seaborn package.

## Acknowledgements

We thank Dr. Gad Getz for valuable discussions of this work. E.R. is supported by funds from MGH, the Broad Institute and the NCI. M.Q. is supported by an MGH ECOR Claflin Award to E.R. A.A.L. is supported by the Mark Foundation for Cancer Research and is a Leukemia & Lymphoma Society Scholar. A.A.L. and E.R. are supported by the Bertarelli Rare Cancer Fund at Harvard Medical School.

## Author contributions

E.R. and A.A.L conceived the study. M.Q., J.P. and E.R. developed computational methods and contributed analysis. I.M. contributed analysis. M.Q. and E.R. wrote the manuscript.

## Supplementary Figure Legends

**Supplementary Figure 1: A,** Y gene expression for TCGA male normal samples ordered by median expression value. **B**, Y gene expression for TCGA male tumor samples ordered by median expression value. **C,** Distribution of Y/housekeeping gene expression ratio in TCGA male and female normal and tumor samples. **D,** Y/housekeeping expression ratio for male tumor samples called as fLOY (blue) and WT (grey).

**Supplementary Figure 2: A,** Bar chart depicts concordance between exome-based LOY calls and expression-based fLOY calls. **B,** Tumor purity for male tumor samples identified as WT by both methods, discordant calls and loss of Y (fLOY/LOY) shows that LOY calls missed by expression have lower purity. **C,** Fraction of tumors by tumor type identified as fLOY (compare to Figure 2A). **D,** Comparison of exome-inferred LOY and expression-based fLOY.

**Supplementary Figure 3: A,** Distribution of *XIST* expression by X alteration in female TCGA tumors. **B,** X alterations in TCGA female tumors by tumor type. Tumor types are ordered by fraction of WT samples. **C**, Distribution of fraction of LOY in male tumors vs fraction of LOX in female tumors.

**Supplementary Figure 4: A,** Fraction of somatic LOY events in tumors from Asian, Black and white participants (only ancestry groups with sufficient sample numbers were included in this analysis). **B,** Distribution of X alterations by race (compare to panel **A**).

**Supplementary Figure 5: A,** Distribution of average Y gene expression among male tumors from Laurent et al.^29^ **B,** Percentage of patients who develop distant metastasis from Laurent et al, by tumor LOY status (compare to **Figure 4C**). **C,** TCGA gene expression for Y-linked genes *KDM5D* and *UTY (KDM6C)* by tumor LOY status. **D,** Different metrics for single-cell quality (# UMIs per cell, # detected genes per cell, log_10_ (genes per unified molecular identifier (UMI)), average housekeeping gene expression level) for WT and LOY cells in uveal melanoma samples shows that LOY is not due to fewer overall genes detected and missed Y genes. UMI: unique molecular identifier. **E,** Fraction of patients with distant metastasis in WT and LOX tumors.

**Supplementary Figure 6:** Differential gene expression between LOY and WT male cell lines from the CCLE.

**Supplementary Figure 7:** Representative examples for male and female sex chromosome copy number calls. Left: male tumors with WT, LOY, pLOY events. Right: WT, LOX, pLOX in female tumors.

## Supplementary Tables

**Supplementary Table 1:** Y chromosome status for male TCGA tumors

**Supplementary Table 2:** X chromosome status for female TCGA tumors

**Supplementary Table 3:** LOY, fLOY and LOX rates by tumor type

**Supplementary Table 4:** GISTIC2 Y arm-level results by tumor type

**Supplementary Table 5:** Cancer gene mutation enrichment in LOY tumors

**Supplementary Table 6:** GISTIC2 focal CNVs on Y in samples without LOY

**Supplementary Table 7:** MutsigCV results for Y-linked genes.

**Supplementary Table 8:** Hazard ratios, *P* and *Q*-values for different tumor types

**Supplementary Table 9:** Differential gene expression analysis in UVM

**Supplementary Table 10:** DepMap dependencies in fLOY and WT cell lines

