## Supplementary Figures for "Loss of chromosome Y in primary tumors"

### Supplementary Figure 1

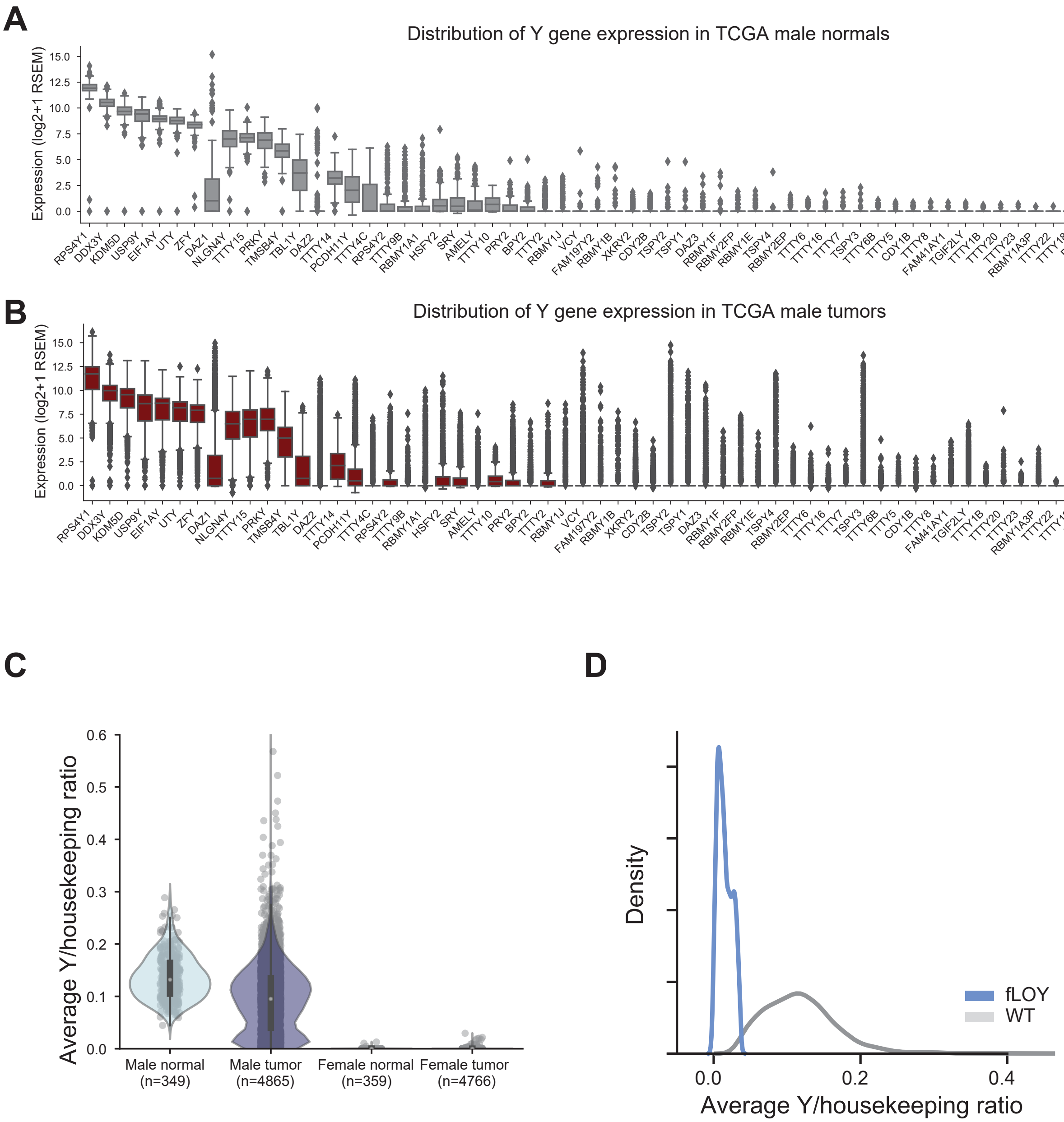

### Supplementary Figure 2

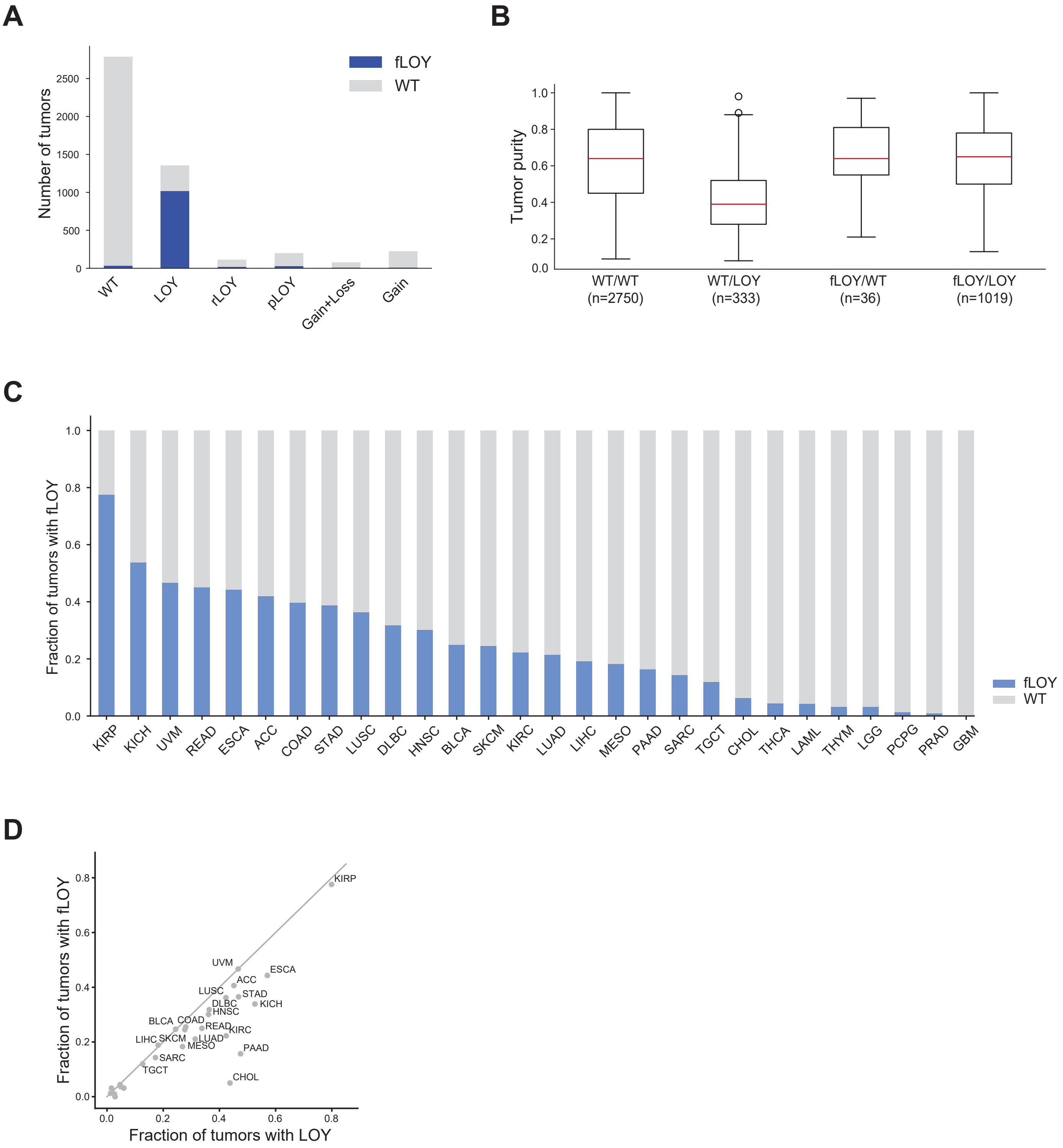

### Supplementary Figure 3

A

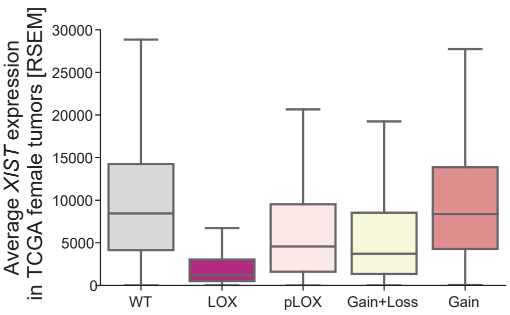


B

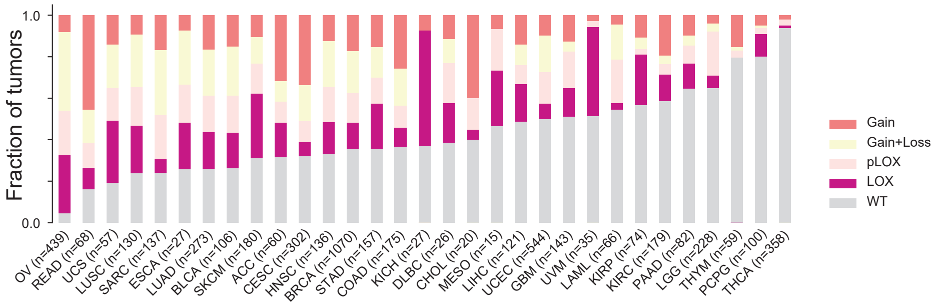


C

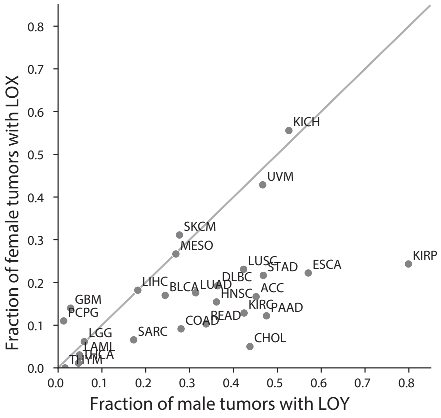

### Supplementary Figure 4

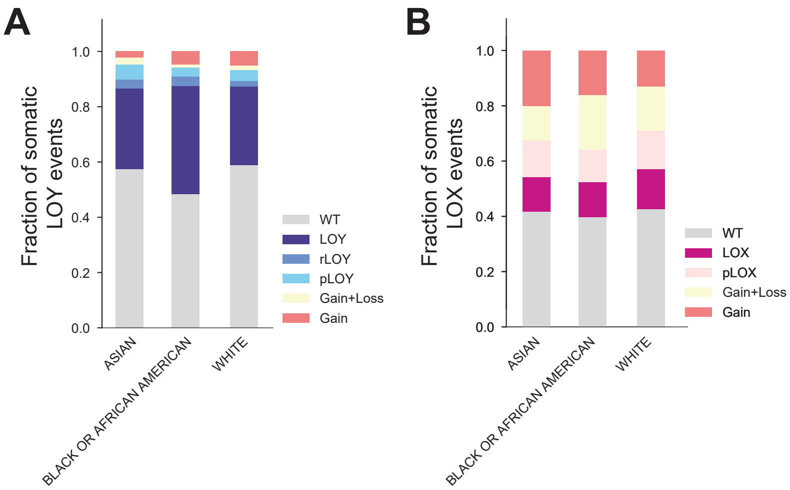

### Supplementary Figure 5

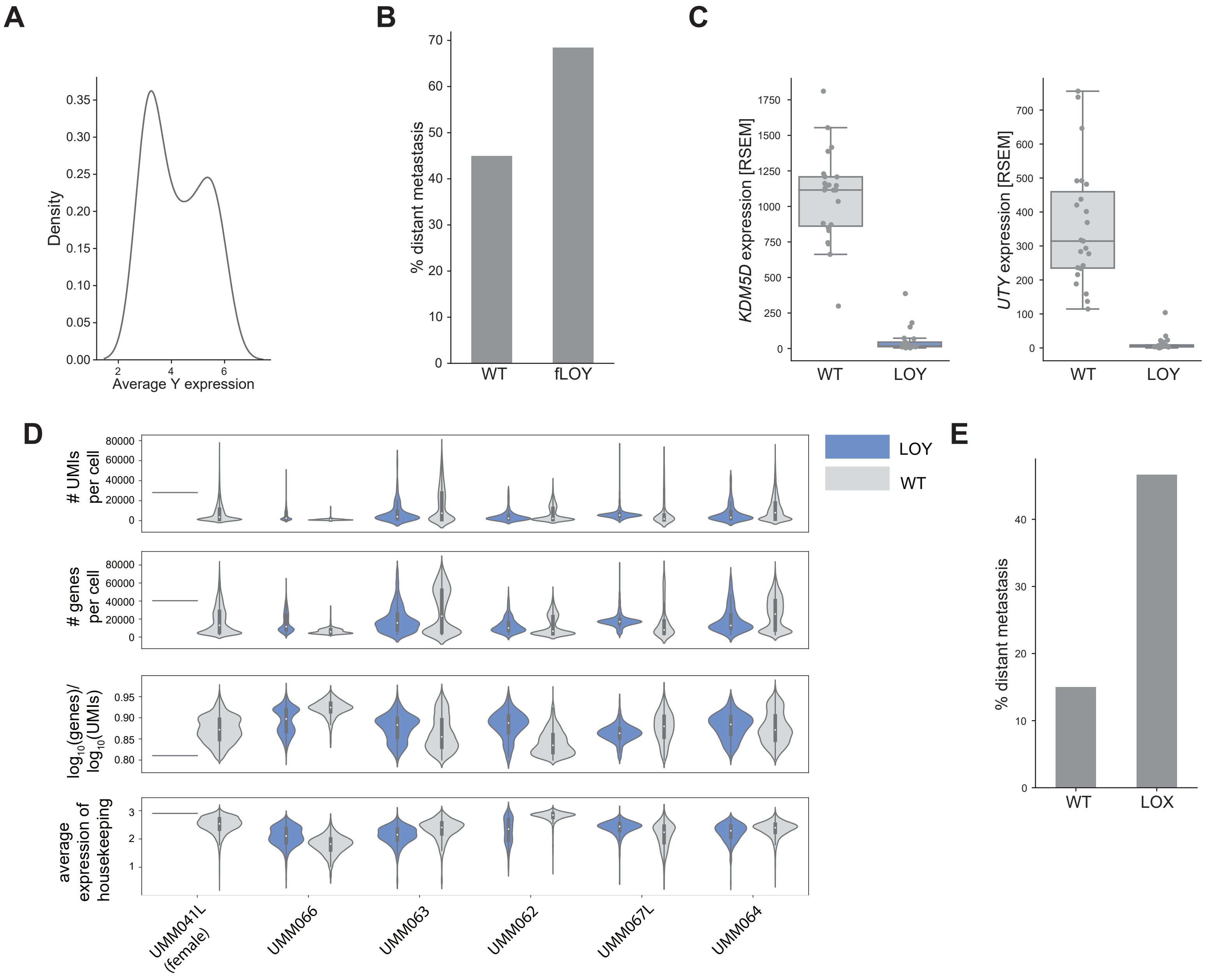

### Supplementary Figure 6

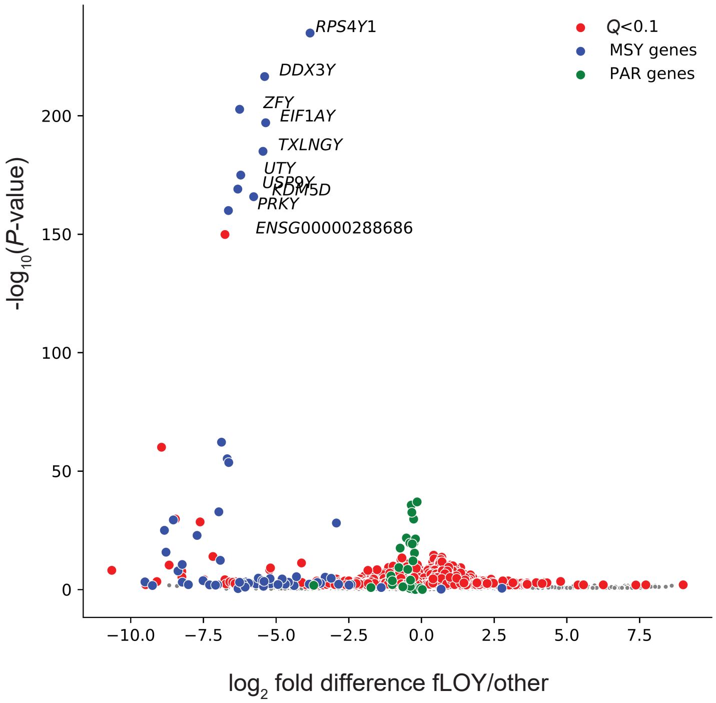

### Supplementary Figure 7

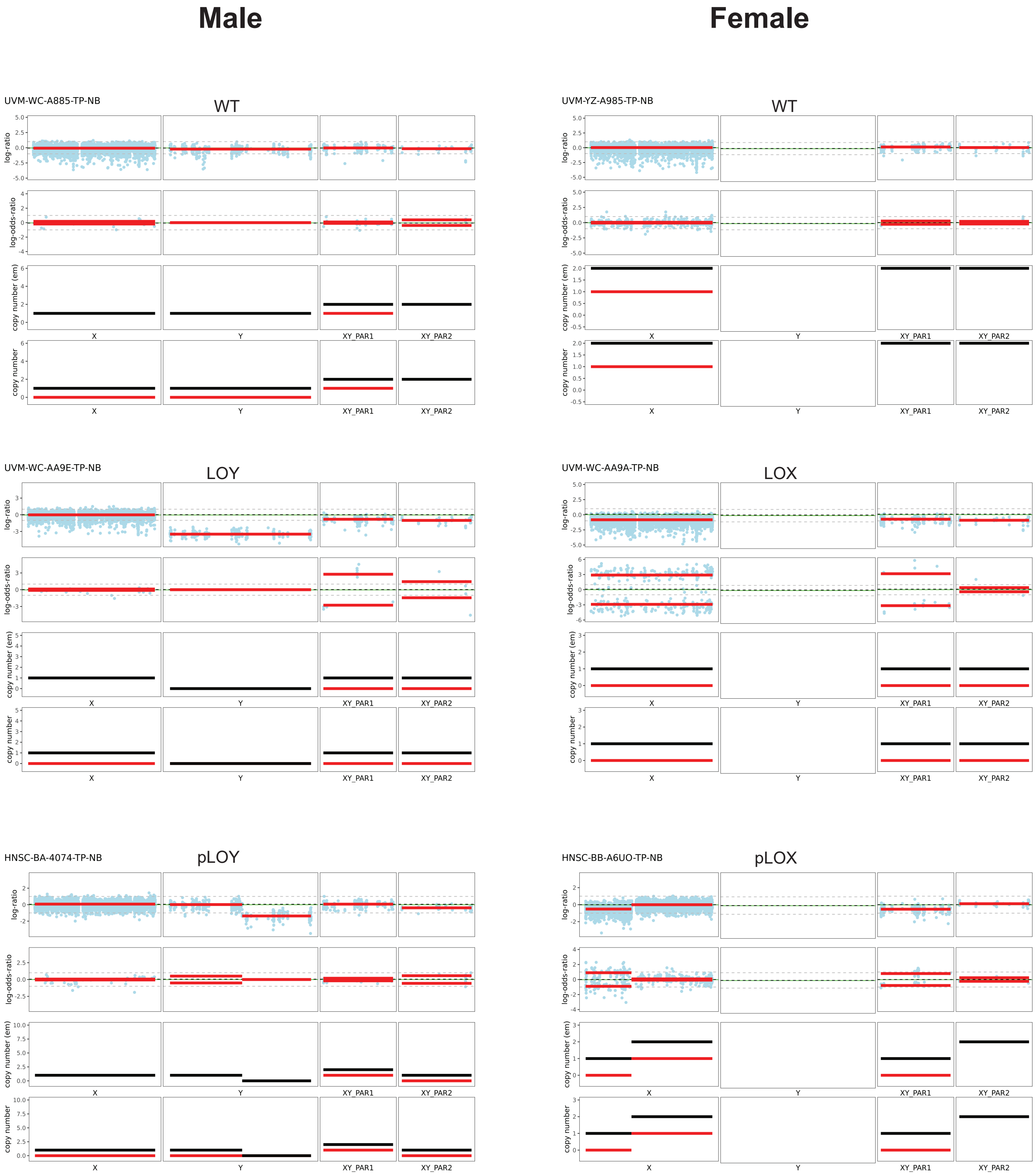
